# Control of lumen geometry and topology by the interplay between pressure and cell proliferation rate in pancreatic organoids

**DOI:** 10.1101/2024.05.29.596462

**Authors:** Byung Ho Lee, Kana Fuji, Heike Petzold, Phil Seymour, Siham Yennek, Coline Schewin, Allison Lewis, Daniel Riveline, Tetsuya Hiraiwa, Masaki Sano, Anne Grapin-Botton

**Affiliations:** Max Planck Institute of Molecular Cell Biology and Genetics, Dresden, Saxony 01307, Germany; Universal Biology Institute, Graduate School of Science, The University of Tokyo, Tokyo, Japan; The Novo Nordisk Foundation Center for Stem Cell Biology, Copenhagen, København N, 2200, Denmark; IGBMC, Université de Strasbourg, CNRS, INSERM, Institut de Génétique et de Biologie Moléculaire et Cellulaire, F-67400 Illkirch, France; Institute of Physics, Academia Sinica, Taipei, Taiwan; Mechanobiology Institute, National University of Singapore, 117411, Singapore; Institute of Natural Sciences, Shanghai Jiao Tong University, Shanghai, China; Cluster of Excellence Physics of Life, TU Dresden, Dresden, 01062, Germany; Paul Langerhans Institute Dresden of the Helmholtz Zentrum München at the University Clinic Carl Gustav Carus of Technische Universität Dresden, Helmholtz Zentrum München, Neuherberg, Germany

## Abstract

Many internal organs in multicellular organisms comprise epithelia which enclose fluid-filled cavities. These are referred to as lumens and their formation is regulated by a wide range of processes, including epithelial polarization, secretion, exocytosis and actomyosin contractility [1, 2]. While these mechanisms have shed light on lumen growth, what controls lumen morphology remains enigmatic. Here we use pancreas organoids to explore how lumens acquire either a spherical shape or a branched topology [3]. Combining computational simulations based on a phase field model with experimental measurements we reveal that lumen morphology arises from the balance between the cell cycle duration and lumen pressure, with more complex lumen at low pressure and fast proliferation rates. Moreover, the manipulation of proliferation and lumen pressure *in silico* and *in vitro* is sufficient to alter and reverse the morphological trajectories of the lumens. Increasing epithelial permeability of spherical lumens lead to lower lumen pressure and converts their morphology to complex lumen shapes, highlighting its crucial role. In summary, the study underscores the importance of balancing cell proliferation, lumen pressure, and epithelial permeability in determining lumen morphology, providing insights relevant to other organs, for tissue engineering and cystic disease understanding and treatment [4].

## Main

Internal organs frequently comprise epithelia that delineate fluid-filled lumens. These lumens vary in shape, ranging from sacs like the bladder to single tubes such as the intestine, or complex networks as seen in the kidney or in many glands including the pancreas [5–7]. Studies exploring lumen formation have leveraged various model systems, including zebrafish, Drosophila and mouse, to understand how these luminal and ductal structures form [8]. These luminal and ductal structures play a pivotal role in organ functionality, serving as essential transport and delivery networks; any dysmorphogenesis in these structures can lead to severe pathological conditions [9].

Mechanistic studies of lumen morphogenesis have focused primarily on the mechanisms of polarity acquisition and lumen growth, notably using cell lines cultured in three dimensions (3D) such as the MDCK (Madin-Darby canine kidney) system [10]. Though this yielded valuable insight, lumen morphology typically presents as a single sphere in these systems. In addition, organoid models, which more closely mimic physiological organs, often feature a single spherical lumen [1, 10–13]. However, they can exhibit more complex geometries and topologies such as outpocketings around a spherical core [14], multiple small lumen [13, 15] or networks of thin tubes [3, 16], whose formation and diversity deserves more mechanistic understanding.

The intricate process of lumen morphogenesis involves multiple cellular mechanisms such as epithelial polarization, secretion, vesicle trafficking and fusion, cortical contractility, as well as cell death and rearrangement [1, 2]. Numerous proteins controlling these processes have been identified. In addition, recent focus on the physics of lumen formation, notably the balance of luminal forces, such as hydrostatic pressure and cell mechanics has provided a more biophysical view of lumenogenesis [12, 14, 15, 17–24]. These studies have largely been limited to spherical lumen.

Pancreatic organoids can form either large spherical lumen or narrow complex interconnected lumen structures, depending on the culture medium [3]. In this study, we investigate how morphological trajectories arise and can be altered or reversed from a fundamental mechanical perspective. Combining experimental insights with multicellular phase field modeling, our assessment of proliferation and lumen pressure as well as targeted interventions unveil how the balance between cell cycle rate and luminal pressure orchestrates the diverse spectrum of lumen morphologies. Our work shows that the leaky epithelium found in organoids prevents high luminal pressure and together with fast proliferation conditions enables the formation of narrow complex interconnected lumen similar to those found *in vivo*.

## Results

### Two distinct morphological lumen trajectories in pancreas organoids

Our previous work has shown that dissociated cells from embryonic day 10.5 (e10.5) pancreatic buds can generate spherical single-lumen and interconnected complex-lumen organoids in different media (hereafter, called spherical organoids and branching organoids for simplicity) (Supplementary Figure 1a) [3]. Specifically, the epithelium of the branching organoids forms a multilayered and branched structure by day 6, whereas the spherical organoids maintain their characteristic epithelium monolayer throughout culture growth (Figure 1a).

**Figure 1.**
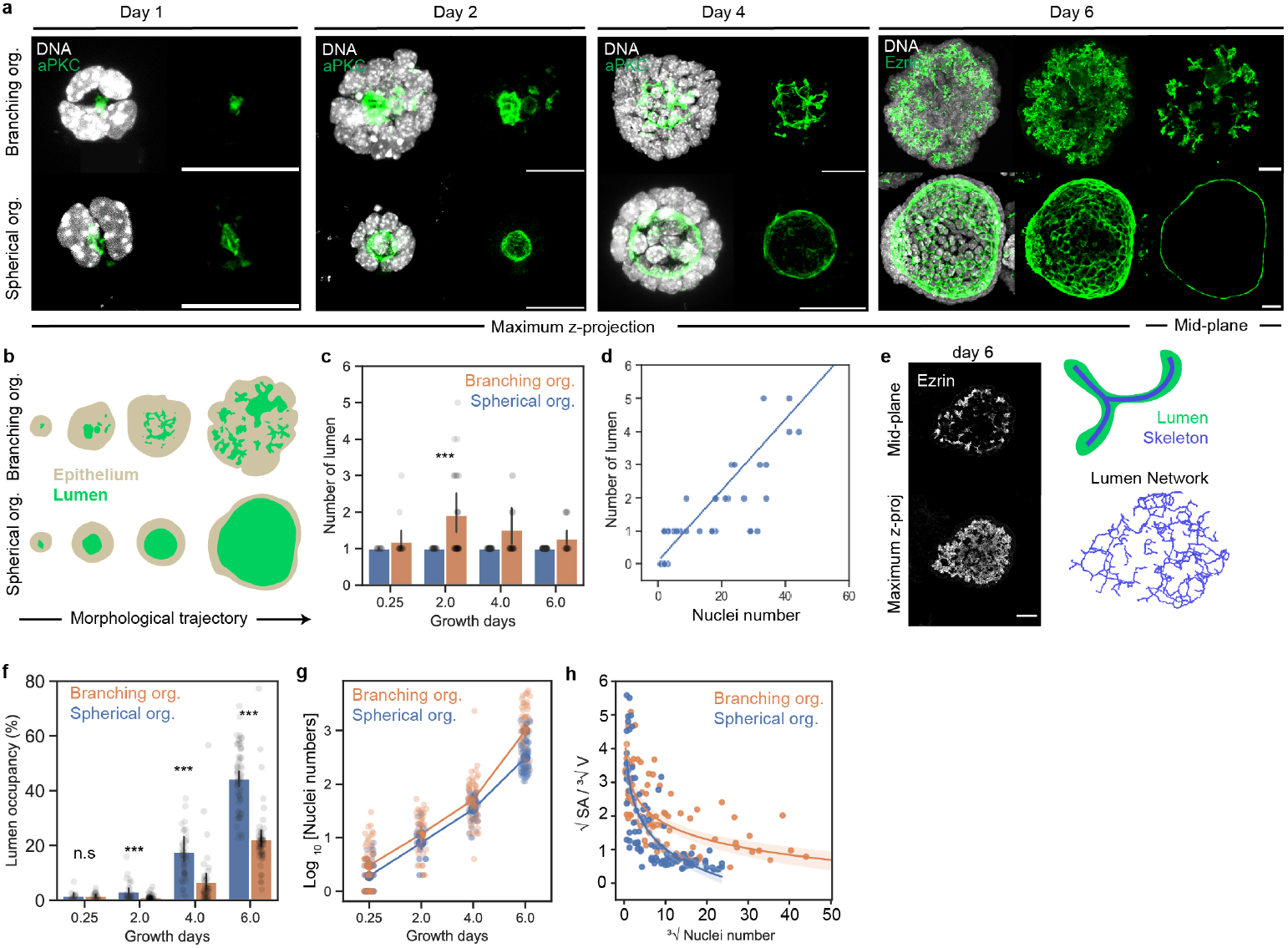
Divergent lumen morphology and topology trajectories in pancreatic branching and spherical organoids. **(a)** Immunofluorescence images of spherical and branching organoids at day 1, day 2, day 4, and day 6 of culture growth in Matrigel. Green: apical marker aPKC (day 1-4) and ezrin (day 6), white: DNA. Scale bar = 20 *µ*m. **(b)** Schematic representing the morphological trajectories of branching and spherical organoids. Beige: epithelium, and green: lumen. **(c)** Number of lumen (quantified in 3D) of spherical and branching organoids at various days of culture growth. **(d)** Relationship between lumen number and cell number of branching organoids at day 2. N=3, n=92 branching organoids. **(e)** Left: Immunofluorescence images showing branching organoids at day 6 with Ezrin marking the lumen. Right: Schematic showing the skeletonization of a lumen (top) to obtain a 3D lumen network (bottom) depicting lumen topology. Scale bar=20 *µ*m. **(f)** Lumen occupancy in percentage (quantified in 3D) of spherical and branching organoids at various days culture growth of culture growth. N = 3-5 exp; n = 220 branching organoids, 194 spherical organoids. Error bar: 95% confidence intervals. **(g)** Quantification of number of nuclei (expressed as log_10_[number of nuclei] of branching and spherical organoids. N=3-5 exp; n=220 branching organoids, 194 spherical organoids. **(h)** Quantification of the relationship between the lumen surface : volume ratio 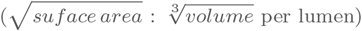 vs number of nuclei 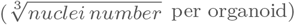. Linear regression estimated of the form lumen surface:volume ratio ∼log(number of nuclei). N = 3-5 exp, n = 165 branching organoids, 194 spherical organoids. We conducted significance using the Mann-Whitney test.

Divergence in the lumen morphology became evident from day 2 onwards, where spherical organoids typically developed and maintained a single lumen throughout the culture growth whereas branching organoids frequently formed multiple lumens (Figure 1b & c). When multiple lumens were present, a larger star-shaped lumen and smaller peripheral lumens were observed. Our previous work has shown that lumens form in two ways in this system, either by cavitation between the small number of cells that seed an lumen or at the time of cell division at the abscission point (Supplementary Figure 1b) [13]. On day 2, the number of lumen formed was proportional to the number of cells in a branching organoid (Figure 1d). Notably, 3D analyses showed that the average number of lumens in the branching organoids peaked on day 2 before decreasing (Figure 1c). This revealed the progressive formation of a hyperconnected network most likely due to the emergence of connections between lumens (Figure 1e).

To further elucidate lumen volume evolution, we analyzed lumen occupancy, which is defined as the 3D lumen volume relative to the total organoid volume (expressed as a percentage in Figure 1f). Both organoid systems showed increasing lumen occupancy over time. Yet from day 2 onward, branching organoids consistently exhibited lower lumen occupancy compared to spherical organoids. Meanwhile, the epithelium of the branching organoids displayed higher number of cells throughout culture (Figure 1g and organoid volumes in Supplementary Figure 1c). Initially, the surface-to-volume ratio of lumens decreased similarly in both systems, consistent with a phase of lumen growth (Figure 1h). As the cell number increased, the branching organoids displayed higher surface to volume ratio than the spherical organoids as expected for a system with multiple small lumens or more convoluted lumens (Figure 1h). In contrast, spherical organoids minimize their surface-to-volume ratio, maintaining their spherical lumen shape. The measurements established here provide a foundation to address what causes differences in both topological (number of lumens) and geometric features (lumen occupancy, surface-to-volume ratios) between spherical and branching organoids.

### Phase field multicellular modelling and experimental validation reveal a major role of lumen pressure and cell cycle duration on the morphological diversity of lumens

Since the increase in cell number was faster in branching organoids, we hypothesized that the creation of new lumen at cell division might drive differences in lumen shape and topology, notably leading to an increase in the number of lumens and apical surface. Additionally, given studies suggesting that lumen growth can be regulated by the pressure difference between the lumen and the exterior environment, we explored whether the balance between lumen pressure and epithelial proliferation rate is a key factor distinguishing these systems [17, 21, 25–29].

To connect single cell dynamics (cell growth and division) with luminal nucleation and growth, we turned to a theoretical investigation by applying phase field modelling in 2D. To mimic organoids, we incorporated a force balance for each cell, considering cortical surface tension, adhesion with neighboring cells, and the neighboring lumen with its osmotic pressure difference to the external environment, *ξ*. As we observed differential growth in branching and spherical organoids, we included the parameter *τ*_*V*_, which captures the time interval between cell divisions in the absence of mechanical constraints. Moreover, the axis of cell division was determined by the force balance regulating the spindle positioning in the dividing cell (see Section 0.13 in Materials and Method for more details and [25, 30]).

In brief, the cells grow in volume (*V*_*i*_) until they reach a target volume (*V*_*target,i*_) which is time-dependent (*t*) and can be tuned by *τ*_*V*_ (Figure 2a i). Moreover, cell division occurs when the target volume (*V*_*d*_) is reached (Figure 2a ii). Therefore, the *τ*_*V*_ parameter provides control of the cell cycle duration *in silico*. Meanwhile, the tuneable parameter to control lumen growth is its osmotic pressure difference to the external environment *ξ*, which is kept constant through numerical simulations and identical in all lumens for each organoid *in silico* (Figure 2a iii; see Section 0.13 in Materials & Methods for more details and [25]). With these rules of cell growth & division and lumen-to-exterior osmotic pressure difference, the *in silico* organoids grow and lumen growth & fusion can be observed (Figure 2b i and Supplementary Video 1). The phase diagram given by the numerical simulations based on this model revealed that both *τ*_*V*_ and *ξ* affected lumen size, shapes and numbers (Figure 2b ii). Multiple areas in the phase diagram displayed lumen features that were comparable to our branching and spherical organoids (branching organoid: orange outline, spherical organoids: blue outline in Figure2b ii & c). Branching organoids were found in regions corresponding to faster proliferation and lower pressure than spherical organoids.

**Figure 2.**
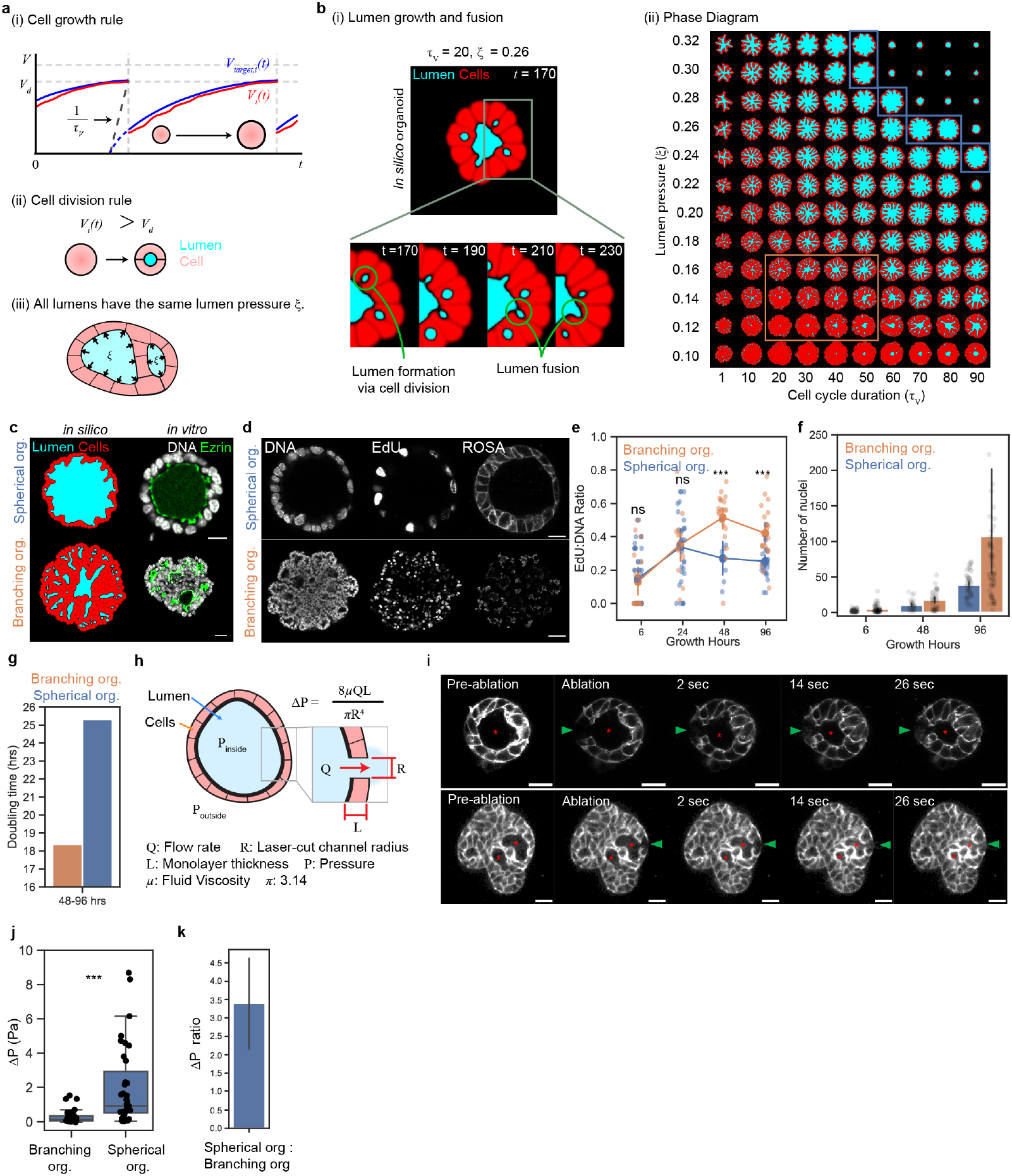
Organoids with complex lumen form at low lumen-to-exterior pressure difference and fast proliferation rates. **(a)** Schematics describing the rules and parameters (*τ*_*V*_ and *ξ*) of the phase field model that governs cell growth (i), cell division (ii), and lumen-to-exterior osmotic pressure (iii). **(b)** *in silico* lumen and organoid growth. i) *in silico* simulation of lumen growth and formation via cell division. ii) Phase diagram capturing the wide spectrum of lumen and organoid morphologies as a function of *ξ* and *τ*_*V*_. **(c)** Qualitative and morphological resemblance between *in vitro* spherical and branching organoids and *in silico* organoids. Scale bar = 20 µm. **(d)** Immunofluorescence images of spherical and branching organoids after 2 hours of EdU labelling. Scale bar= 10 µm (top row) and 30 µm (bottom row). **(e)** EdU:DNA ratio of spherical and branching organoids at various hours of culture growth. N = 2-5 exp; n = 72 branching organoids, 109 spherical organoids. Error bar: 95% confidence intervals. **(f)** Nuclei counts per organoid at various hours of culture growth. Error bar: 95% confidence intervals. N = 3 exp; n = 180 branching organoids, 129 spherical organoids. **(g)** Quantification of doubling time for branching and spherical organoids from 48 hours to 96 hours of culture growth. **(h)** Schematic of the laser-ablated organoid and measurements used in the Hagen-Poiseuille equation to infer lumen-to-exterior hydrostatic pressure difference. **(i)** Laser ablation experiment montage on a branching and spherical organoid expressing ROSAmTmG. Red asterisk: targeted lumen (connected lumen in 3D for branching organoid), green arrowhead: channel created by laser ablation. Scale bar = 20 µm. **(j)** Inferences of lumen-to-exterior hydrostatic pressure difference ΔP of the branching and spherical organoid lumens on day 6 of culture. N = 3 exp; n = 36 branching organoids, 34 spherical organoids. **(k)** Quantification of spherical organoid: branching organoids lumen-to-exterior hydrostatic pressure difference ΔP ratio. Error bar: standard error of the mean over average ratios (See Supplementary Figure 3f). We conducted significance using the Mann-Whitney test..

To experimentally test these observations and validate the model, we sought to characterize the difference in lumen-to-exterior hydrostatic pressure and cell cycle duration of both organoid systems. To address whether the branching organoids proliferate faster, we performed an EdU (5-ethynyl 2’
s - deoxyuridine) incorporation assay, labelling cells in S-phase within the organoids. We treated the organoids with EdU for 2 hours prior to fixation at 6, 24, 48, and 96 hours of culture growth (Figure 2d). To analyze the difference in proliferation, we quantified the ratio between cells that were EdU positive (S-phase) and the total number of nuclei. We observed that at 6 hours and 24 hours both branching and spherical organoids displayed comparable EdU:DNA ratios. However after 24 hours the spherical organoids exhibited a consistently lower relative number of EdU positive cells compared to the branching organoids (Figure 2e). These experiments showed that both organoid systems increase their potential to proliferate and start diverging from day 2 onwards. This observation coincides with the similar lumen morphologies the two organoids share at day 2 (Figure 1a & b). Since the interval between cell divisions *τ*_*V*_ was found to be an important control parameter in the numerical simulations, we calculated the doubling time for both organoid systems by utilizing the average cell number in organoids at later culture days (Figure 2f), *i.e*., when the average morphologies of the branching and spherical organoids were different (day 2 to 4) (Figure 1a & b). Assuming exponential growth and neglecting contribution from cell death, we found that the doubling time of the branching organoids was 1.4 faster than that of the spherical organoids from 48 to 96 hours (Figure 2g).

Another control parameter which played a key role in the phase diagram was *ξ*. We asked whether the branching and spherical organoids exhibited distinct lumen-to-exterior osmotic pressure difference. Through mathematical modeling using the phase field model, we found that the lumen-to-exterior osmotic pressure shares a linear relationship with hydrostatic pressures (Supplementary Figure 4: see Section 0.13.2 in Material & Methods). With this confirmation, we performed laser ablations to infer lumen-to-exterior hydrostatic pressure differences using the Hagen-Poiseuille equation [12, 21, 29, 31]. Laser ablation creates a conduit across the epithelium layer between the lumen and the external environment (Figure 2i). Upon laser ablation, we observed a decrease in the lumen volume (measured in 3D) and fluid expelled from the lumen indicating a higher lumen-to-exterior hydrostatic pressure than around the organoids (Supplementary Figure 3a - c). To calculate the hydrostatic pressure difference (ΔP), we quantified the flow rate across the conduit based on lumen volume change, the conduit and epithelium length and diameter (Figure 2h) [12]. To estimate lumen fluid viscosity, we segmented and tracked Cell-Mask positive particles in the lumen to calculate their 3D mean squared displacement. Using these curves, we derived the diffusion coefficient and then estimated the lumen’s viscosity via the Stokes-Einstein equation (Supplementary Figure 2a - g). With this approach, we estimated that the mean lumen fluid viscosity was 2.06±0.35 mPa·s (Supplementary Figure 2h - i). As the lumens in branching organoids were too narrow to make such measurements, we assumed that the lumen viscosity in both organoid systems was similar and applied the average lumen viscosity obtained from the spherical organoids to infer ΔP of both systems. With the lumen viscosity, flow rate, and conduit dimensions created by the laser cut, we obtained mean ΔP of 0.28 Pa for the branching organoids and 1.46 Pa for the spherical organoids on day 6 of culture (Figure 2j).

Moreover, we observed that the ΔP of the lumen spherical organoids increased with increasing lumen volume. Notably, this was not the case for the branching organoids (Supplementary Figure 3e). We thus compared lumen of the same size for branching and spherical organoids after binning lumen volumes (Supplementary Figure 3f & g). This analysis revealed that ΔP in size-matched spherical organoids is approximately 3.4 times higher than in branching organoids (Figure 2k).

From the linear relationship between ΔP and lumen-to-exterior osmotic pressure (Supplementary Figure 4: see Section 0.13.2 in Material & Methods) we infer that spherical organoids have a 1.9 times higher lumen-to-exterior osmotic pressure than branching organoids. This difference between the two organoids allows to place the *in vitro* branching organoids within the 0.12 to 0.16 *ξ* range, while spherical organoids are in the 0.28 to 0.32 *ξ* range *in silico* (Figure 2b ii). At these values in the Y axis (of the phase diagram), qualitative and quantitative approximations place branched organoid types around values of *τ*_*V*_ = 40 and spherical organoids 1.4 higher at approximately 60. Overall, these results indicate that organoids with faster proliferation and lower lumen pressure align with phase diagram regions featuring more star-shaped lumen geometries and multi-lumen topologies while the opposing organoid features results in a lumen that minimizes its surface to volume ratio to obtain a spherical morphology.

### Branching organoids relax to spherical organoids upon proliferation arrest

Given that our quantitative comparisons for cell proliferation rates and differential lumen-to-exterior pressure involved organoids grown in different culture media, we first aimed to specifically perturb cell proliferation in the same organoid medium. To test the model predictions, we used aphidicolin which slows down or stops proliferation by inhibiting DNA Polymerase-A thus arresting cells in the S-phase of the cell cycle. Lumen morphogenesis is a slow process (time scales of days). Accordingly, short-term (10 hours) aphidicolin treatment was observed to have no impact on intestinal organoid morphologies [14]. Thus we treated the most proliferative of our organoid types, the branching organoids, with DMSO and aphidicolin at day 4, a stage at which they had multiple narrow lumens, and analyzed them at day 6 (Figure 3a). Under these treatment conditions, the cell cycle was arrested leading to no or few organoids with pH3s10 (phospho Histone-3 Serine-10) positive cells (Supplementary Figure 5a). We were unable to identify a dose of aphidicolin that slowed proliferation. By day 6, aphidicolin-treated branching organoids exhibited a more spherical shape, fewer lumen and higher lumen occupancy compared to DMSO-treated branching organoids (Figure 3b & c). These results indicate that stopping proliferation has a drastic impact on lumen morphology in the branching organoids. To mimic proliferation arrest in the *in silico* model, we conducted new simulations at the determined value of *ξ* = 0.12 and *τ*_*V*_ = 40 where cell cycles were halted once the organoids reached a cell number comparable to the average quantified in 2D on day 4, prior to aphidicolin treatment (Figure 3d & e and Supplementary Figure 5c). Moreover, we compared experimental organoids with numerical model phenotypes. Under these conditions, the *in silico* organoids evolved into spheres with single large spherical lumens mirroring the lumen occupancy and numbers (Figure 3f-g, Supplementary Video 2 & 3) observed in experiments (Figure 3h & i). This model shows that proliferation arrest leads to fusions of lumens over time and evolution to a single spherical lumen (Figure 3d & e and Supplementary Figure 5b). The fast proliferation of branching organoids is thus crucial to keep the system out of equilibrium and prevent relaxation of lumens to a single lumen via a slow fusion process.

**Figure 3.**
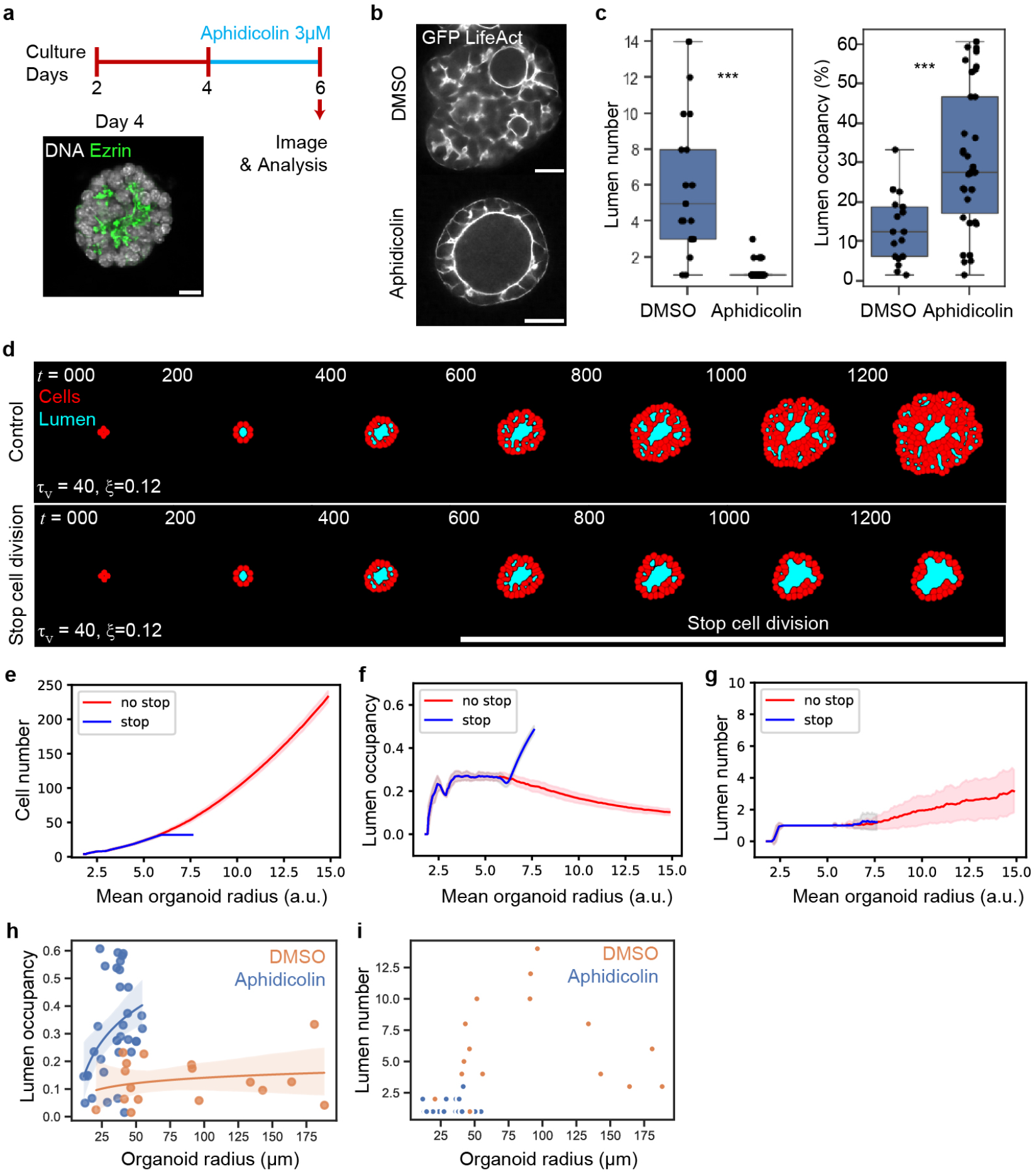
Cell cycle interference reduces the complexity of lumen. **(a)** Schematic showing experimental design of aphidicolin assay. Scale bar = 10 µm.**(b)** Immunofluorescence image of DMSO and aphidicolin treated branching organoids at day 6. Scale bar = 20 µm. **(c)** 2D Quantification of lumen number (left) and lumen occupancy (right; in percentage) of DMSO and aphidicolin-treated branching organoids at day 6. N = 3 exp, n = 17 DMSO, 34 aphidicolin. **(d)** *in silico* simulations of organoids under *τ*_*V*_ = 40, and *ξ* = 0.12. Top row represents a simulation run without stopping cell division. Bottom row represents a simulation run with cell division stopped at 32 cells. Red: cells, blue: lumen. **(e)** Quantification showing the number of cells vs mean organoid radius during the simulation. Red: control with no stop in cell division, blue: stopped cell division at 32 cells. N=50, bold line represents moving average with standard deviation error bars. **(f)** Quantification showing the lumen occupancy vs mean organoid radius during the simulation. Red: control with no stop in cell division, blue: stop cell division at 32 cells. N=50, bold line represents moving average with standard deviation error bars. **(g)** 2D Quantification showing the lumen numbers vs mean organoid radius during the simulation. Red: control with no stop in cell division, blue: stop cell division at 32 cells. N=50, bold line represents moving average with standard deviation error bars. **(h)** 2D Quantification showing the lumen occupancy vs organoid radius of branching organoids under DMSO and aphidicolin. Linear regression estimated of the form lumen occupancy ∼ log(organoid radius). Errobar = 95% confidence interval. N=3 exp, n=34 DMSO, 17 Aphidicolin. **(i)** Quantification showing the lumen number vs organoid radius of organoids under DMSO and Aphidicolin. N=3 exp, n=34 DMSO, 17 Aphidicolin. We conducted significance using the Mann-Whitney test.

### Controlling organoid lumen morphology through manipulation of osmotic pressure

Numerous studies theoretically predict or underscore lumen pressure’s influential role in determining organoid morphologies [12, 21, 25]. However, direct perturbation of lumen osmotic pressure was rarely combined with measurements of the effect on hydrostatic pressure and an assessment of the impact on complex lumen morphologies.

To perturb lumen-to-exterior osmotic pressure, we treated the branching organoids with forskolin, an activator of cystic fibrosis transmembrane regulator (CFTR) ion channels which triggers the secretion of chloride ions and bicarbonate into the lumen. Secretion of ions is expected to increase lumen-to-exterior osmotic pressure [12, 21]. Forskolin treatment from day 4 to day 6 in our branching organoids resulted in inflated lumen and increased lumen occupancy but did not yield a single spherical lumen (Supplementary Figure 6a - d). Measurements of lumen-to-exterior hydrostatic pressure difference at day 6 revealed no significant difference between branching organoids treated with DMSO and those treated with forskolin (Supplementary Figure 6f).

While chloride ion secretion may transiently increase lumen-to-exterior osmotic pressure in the lumen and subsequent water influx, we conclude that it results in a change of lumen volume but does not lead to a sustained increase in lumen-to-exterior hydrostatic pressure in branching organoids (Supplementary Figure 6g). Given the unexpected lack of lumen pressure increase by forskolin, we hypothesized that the epithelium of organoids may not retain solutes and water sufficiently to enable pressure increase.

### Epithelial permeability contributes to the regulation of lumen morphology

Epithelial paracellular permeability or “tightness” regulates the epithelial barrier function against ions, solutes, molecules [32]. To test permeability we added 10kDa fluorescent Dextran-647 into the culture mediums. Comparable to reports of MDCK and certain intestinal organoids, the spherical organoids displayed no or low signals of 10kDa Dextran-647 in the lumen after 3 hours, indicating their “tight” epithelial monolayer [12]. However, branching organoids displayed lumens with 10kDa Dextran-647 signals as well as signal in paracellular spaces indicating higher permeability (Figure 4a). These experiments thus reveal a striking difference in epithelial permeability between branching and spherical organoids.

**Figure 4.**
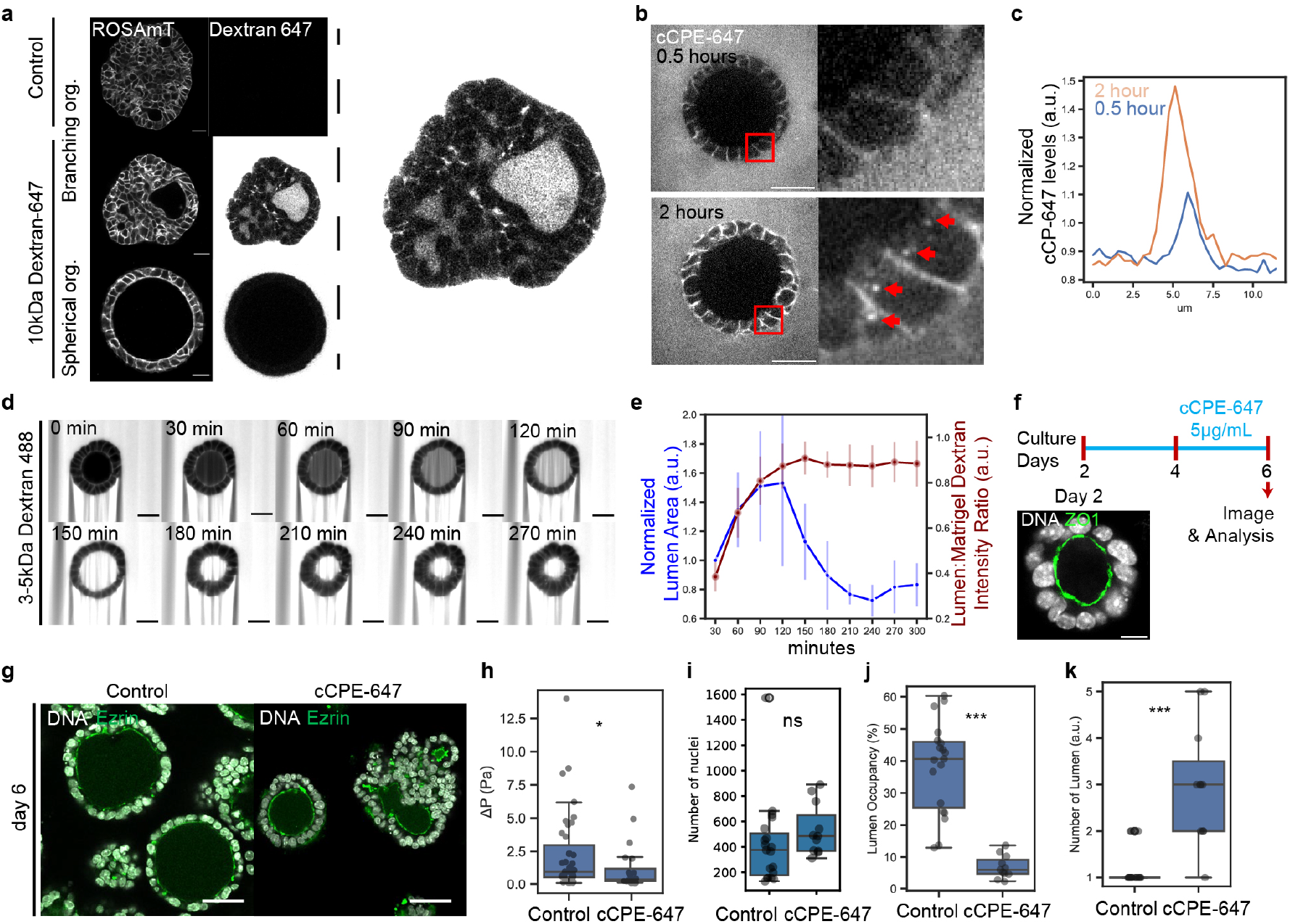
Transformation of spherical organoids into branching organoids through induced permeation. **(a)** Left: Live images of control and 10kDa Dextran-647 treated live branching and spherical organoids 3 hours post-treatment. Right: Enlarged image of branching organoids with 10kDa Dextran-647 signal detected in paracellular spaces and lumens. Scale bars = 20 µm. **(b)** Live images of day 4 spherical organoids treated with cCPE-647 at 0.5 hours and 2 hours of incubation. Fluorescent signals are cCPE-647 in the Matrigel and paracellular spaces. Red arrowheads indicate cCPE-647 puncta in the cytoplasm. Scale bar = 20 µm. **(c)** Line-profile quantification of cCPE-647 (normalized to signals in the Matrigel) across the paracellular spaces at various time points of treatment. **(d)** Montage of live spherical organoids co-treated with Dextran 3-5 kDa-488 and cCPE-647 at various timepoints of treatment. Scale bar = 20 µm. **(e)** Quantification of normalized levels 3-5kDa Dextran-488 in the lumen (mean lumen: mean Matrigel intensity ratios of Dextran 3-5 kDa-488) and mean normalized 2D area of lumen (normalized to lumen area at first time point of image acquisition) at various time points of cCPE-647 treatment. n = 5 spherical organoids. Error bar: standard deviation. **(f)** Schematic showing experimental design of cCPE-647 treatment. Scale bar = 10 µm. **(g)** Immunofluorescence images control and cCPE-647-treated spherical organoids on day 6. Scale bar = 30 µm. **(h)** Inference of lumen-to-exterior hydrostatic pressure ΔP of the control and cCPE-647 treated spherical organoid lumens on day 6 of culture. N = 2-3 exp; n = 39 control, 20 cCPE-647. **(i-k)** 3D quantification of nuclei counts, lumen occupancy, and number of lumens of control and cCPE-647 treated spherical organoids on day 6. N = 2-3 exp; n = 27 control, 17 cCPE-647. We conducted significance using the Mann-Whitney test.

To investigate whether increased permeability leads to decreased pressure and a subsequent change in lumen morphology, we used Clostridium perfringens enterotoxin (CPE) which forms pores in the epithelium and disrupts tight junctions by binding to claudins [33, 34]. Reports show that CPE has the high affinity towards Claudin-3 and -4, and both organoid systems express prominent levels of the two claudins based on transcriptomics (Supplementary Figure 7a) [34, 35]. In addition, non-cytotoxic recombinant forms of CPE (cCPE) have been utilized to manipulate barrier functions via tight junction modulations [34, 35]. Therefore, we synthesized cCPE labelled with ATTO647 maleimide and used it to manipulate permeability and investigate alterations in lumen morphology in the spherical organoids (Supplementary Figure 7d). Within 2 hours of cCPE-647 treatment, signals were detected in the paracellular spaces of spherical organoids followed by cCPE-647 puncta in the cytoplasm, consistent with previous reports of internalization with claudin targets [36, 37] (Figure 4b & c). We then co-treated spherical organoids with 3-5 kDa Dextran-488 and cCPE-647 to assess permeability and monitor alterations in the lumen. Approximately 1 hour after cCPE-647 treatment, we observed an increase in the level of 3-5 kDa Dextran-488 in the lumen, coinciding with an increase in the lumen area (Figure 4d & e). After approximately 2 hours post-treatment, when the 3-5 kDa Dextran-488 levels in the lumen plateaued, we observed a shrinkage in the lumen area. Using laser-ablation, we observed that cCPE-647 lowered the lumen-to-exterior hydrostatic pressure (mean 1.15Pa) compared to the control condition (mean 2.22Pa) (Figure 4h). Moreover, the positive correlation between the ΔP and lumen volume observed in spherical organoids was lost under cCPE-647 treatment (Supplementary Figure 7g).

To evaluate the long-term effects of induced-permeabilization and reduced pressure on lumen morphogenesis, we treated spherical organoids in spherical organoid-medium with cCPE-647 from day 2 to day 6 (Figure 4f) and observed 50% of organoids adopting a branching morphology, exhibiting lower lumen occupancy and higher lumen number (Figure 4j - k & Supplementary Figure 7e). No change in cell number was observed (Figure 4i). Moreover, overall shape of the organoid shell was more branched, as shown by a decreased sphericity (Supplementary Figure 7h).

Collectively, these experiments indicate that permeability has a significant role in regulating lumen pressure with consequences on lumen morphology, independently of cellular proliferation.

## Discussion

While previous work has emphasized the crucial role of apical surface growth via vesicle fusion in the formation and enlargement of lumens in the pancreas and other 3D models, our study reveals another critical factor shaping pancreatic organoids: the permeability of the pancreatic epithelium and its role in maintaining a moderate lumen pressure (Supplementary Figure 3e, 6g, and 7g) [38–42]. Computational models of lumenogenesis have shown that hydrostatic pressure resulting from the equilibrium between osmotic pressure, fluid influx, and paracellular leaks contributes to lumen growth and homeostasis [17–19, 21, 24]. However, pressure has been measured in few biological lumens [12, 22, 26, 28, 29, 43]. Reported heterogeneities in pressure appear to be significantly influenced by specific tight junction components that regulate tension and leakiness [12, 20]. An imbalance between high junctional tension and low lumen pressure has been shown to affect lumen shape in ZO-1/2 knockout MDCK cysts, leading to invaginations in the apical membrane of cells lining the lumen. In contrast, Claudin 1-5 knockouts exhibited no such changes in lumen morphology and no alterations in pressure [12, 20]. Importantly, our experiments with cCPE, a claudin-targeting toxin, substantiate that altering lumen permeability can significantly change hydrostatic pressure and consequently lumen morphology. Different claudins mediate distinct permeability characteristics, as observed in various intestinal organoid models with different tight junction compositions showing differential permeability to Dextran [12, 20, 44]. In pancreas organoids, transcriptomic investigation shows the expression of many claudins as well as major differences in Claudin 4, 10, and 8 between the spherical and branching organoids (Supplementary Figure 7a - c). Claudin 4 expression, one of the main targets of cCPE, is lower in branching organoids than spherical organoids. Claudin 4 has an important role in permeability, in combination with other Claudins (e.g. Claudin 2 and 15) [35]. It is therefore likely that the higher permeability and lower lumen pressure of branching organoids is due to Claudin 4 and that targeting of Claudin 4 by cCPE in spherical organoids reduces their pressure, rewiring them between extreme trajectories from single spherical lumen to multi-lumen [45, 46]. Tissue specificities in the balance of mechanisms controlling pressure (permeability to specific solutes and the rate at which they are secreted in the lumen), apical growth and junctional tension would require further investigations in different organs and organoid proxys.

Another significant parameter influencing lumen shape is the cell division rate, which dictates the rate at which new lumen form at abscission points. Cerruti *et al*., have used cell packing patterns and modelling to evaluate how far from mechanical equilibrium MDCK cysts are. They further showed that this depends on their cell division rate and their cell rearrangement rate, with the latter being at a longer timescale than duplication [47]. While their experimental inhibition of cell division did not reveal drastic changes in lumen shape or topology, this difference can be attributed to the higher pressure regime in MDCK cysts, promoting more spherical morphologies, potentially masking the effects of cell proliferation [12]. In contrast, our study shows that in pancreatic organoids, cell division rate significantly impacts lumen shape. We hypothesize that in branching organoids, lumen volume cannot expand quickly enough to match the increasing apical surface area provided by rapid cell cycles, resulting in multiple small lumens rather than a single large lumen.

Collectively, this study highlights the critical influence of the culture medium composition on modulating physical parameters regulating the morphological trajectories of pancreatic organoids. The high levels of FGFs and EGFs in the organoid medium promote fast cell division [3]. We propose that rapid lumen creation is only partially compensated by slow cell rearrangements enabling lumen fusion. Fast division may potentially contribute to the leakiness of the system, as cells must continually reorganize their junctions.

Our study employs a reductionist 3D culture model using primary cells freshly isolated from *in-vivo* tissues, enabling experiments that would be challenging to perform *in-vivo*. Despite its simplicity, this model system reveals that a finely-tuned balance between cell proliferation, lumen pressure, and epithelial permeability dictates the morphological diversity observed in pancreatic organoids. It reveals mechanisms that are potentially relevant to other organs exhibiting narrow interconnected ducts and to common cystic diseases affecting the pancreas (cystic fibrosis, cysts in pancreatic cancers and pancreatitis) as well as various branched organs. The system could for example be used to test the effect of drugs reverting disease phenotypes for possible therapeutic interventions. Moreover, it paves the way for future research to engineer organ shape in regenerative medicine [4].

## Materials and Methods

### 0.1 Animals and permit

All experiments were performed in accordance with the German Animal Welfare Legislation (“Tier-schutzgesetz”) after approval by the federal state authority Landesdirektion Sachsen (license DD24.1-5131/451/8). Mice were kept in standardized specific-pathogen-free (SPF) conditions at the Biomedical Services Facility (BMS) of Max Planck Institute of Molecular Cell Biology and Genetics. Genetically modified mouse lines LifeAct-EGFP [48] and ROSAmT/mG [49].

### 0.2 Pancreatic organoid culture

Mouse embryonic stage e10.5 was defined as noon of the day when the vaginal plug was detected in the mother. Pancreatic buds were dissected from e10.5 mice and mesenchymal cells were removed using Tungsten needles [50]. To obtain cell aggregates, the buds were dissociated using TrypLE (12604013, ThermoFisher Scientific) treatment for 12 min in a 37°C incubator, followed by mechanical dissociation using pulled glass capillaries (BR708707, BRAND/Merck). The cell aggregates were seeded into 75% Matrigel (356231, Corning) in 8 well glass bottom plates (80826, Ibidi) and left to polymerize at 37°C in an incubator for 10 min. To grow branching organoids, a medium composed of 25 ng/mL murine-FGF1 (450-33, Perprotech), 25 ng/mL murine-EGF (315-09, Perprotech), 2.5 U/mL Heparin (7980, Stemcell), 10 *µ*M Y-27632 -dihydrochloride ROCK inhibitor (Y0503, Sigma Aldrich), 16 nM Phorbol-12-myristat-13-acetate (524400, Milipore), 100 ng/mL murine-FGF10(450-61, Perprotech), 500 ng/mL murine-Spondin-1 (315-32, Perprotech), 10% Knockout serum (10828-028, Gibco), 1% Penicillin-Streptomycin (15140-122, Sigma Aldrich) and DMEM/F12 (1:1) 1x (+) L-Glutamine (11320-033, Gibco) was added. To grow spherical organoids, a medium composed of 64 ng/mL murine-FGF2 (450-33, Peprotech), 10% B27 supplement (17504-044, Gibco), 10 *µ*M Y-27632 -dihydrochloride ROCK inhibitor (Y0503, Sigma Aldrich), 1% Penicillin-Streptomycin (15140-122, Sigma Aldrich), and DMEM / F12 (1:1) 1x (+) L-Glutamine (11320-033, Gibco) was added. The organoids were grown for 6 days in culture in an incubator at 37°C 5% CO_2_. Medium exchange was carried out every 2 days.

### 0.3 Pharmacological & Chemical treatment

To stop proliferation, pancreatic organoids were treated with aphidicolin (32774, Cell Signaling Technology/Merck), a DNA Polymerase A inhibitor. Aphidicolin was dissolved in DMSO (800153, Cell Signaling Technology) and a final concentration of 3 µM concentration were used for 2 days. To inflate the lumen, the CFTR activator forskolin (32774, Tocris Bioscience) at were used 10 µM for 2 days.

### 0.4 Immunofluorescence

Pancreatic organoids were fixed with 4% Formaldehyde (28908, ThermoFisher Scientific) in PBS for 30 min at room temperature. Samples were blocked and permeabilized in 0.25% Triton (T8787, Sigma Aldrich), 1% bovine serum albumin (BSA; A3059, Sigma Aldrich) in PBS for 6 h at room temperature. Pancreatic organoids were incubated in primary antibody solution in 0.25% Triton, 1% BSA in PBS overnight at 4°C, and in secondary antibody solution in 0.25% Triton, 1% BSA in PBS overnight at 4°C. To stain nuclei, Hoechst solution (34580, Invitrogen) or DAPI (ab228549, Abcam) was added at 1:1000 in 0.25% Triton, 1% BSA in PBS for 4 hours at room temperature after the incubation of secondary antibody.

The primary antibodies used to mark the lumen were anti-Aurora B (Becton Dickinson), anti-Ezrin (3C12) (sc-58758, Santa Cruz), anti-Mucin-1 (MH1(CT2)) (MA5-11202, Thermofisher Scientific), anti-ZO1(1A12) (339100, ThermoFisher Scientific), anti-aPKC (H1) (sc-17781, Santa Cruz), and Alex-488 Phalloidin (A12379, Thermofisher Scientific). To mark epithelial cells anti-Ecad (M108, Takara Bio), and anti-Sox9 (AB5535, Merk) were used. To mark proliferating cells, anti-phosho Histone-3 Serine-10 (3H10) (05-806, Millipore/Sigma Aldrich) was used. Secondary antibodies used were goat anti-Armenian hamster IgG H&L (Alexa Fluor 568) (ab175716, Abcam), goat anti-Mouse IgG H&L (Alexa Fluor 488) preadsorbed (ab150117, Abcam), anti-Mouse IgG H&L (Alexa Fluor 647) preadsorbed (ab150111, Abcam). Primary and secondary antibodies were used with a dilution factor of 1:400 for all immunofluorescence experiments.

### 0.5 Clostridium Perfringens enterotoxin (CPE) fragment expression, purification and labelling

#### 0.5.1 Cloning

The non cytotoxic, claudin binding, C-terminal domain of CPE (Supplementary Figure 7d) was been expressed and purified according to Tachibana *et al*. with few modifications [51]. In details, codon optimized cCPE fragment, amino acids 184-319, sequence: ERCVLTVPSTDIEKEILDLAAATERLNLT-DALNSN PAGNLYDWRSSNSYPWTQKLNLHLTITATGQKYRI LASKIVDFNIYSNNFNNLVKLE-QSLGDGVKDHYVD ISLDAGQYVLVMKA NSSYSGNYPYSILFQKF (Twist Biosciences) was cloned into p7XNH3 vector and tagged with a N-terminal 10xHis tag cleavable with human rhinovirus 3C protease [52].

#### 0.5.2 Expression

E.coli T7 Express strain (New Englad Biolabs) was transformed with p7XNH3-10xHis-3C-cCPE and pRare plasmids. A preculture was grown in LB media supplemented with 1% glucose, 30 *µ*g/ml kanamycin (kan) and 17 *µ*g/ml cloramphenicol (cm), overnight at 37°C, shaking at 150 rpm (Kunher shaker). E.coli cultures for induction were grown in Terrific broth (TB), supplemented with 90 *µ*g/ml kanamycin and 17 *µ*g/ml chlorampphenicol antibiotics at 37°C. When the optical density at 600 nm reached a value of 0.6-0.8, cultures were moved into a 18°C shaking incubator. Protein expression was induced with 0.2 mM IPTG (Sigma) final concentration, overnight at 18°C.

#### 0.5.3 Lysis, IMAC, His tag removal and Size Exclusion Purification Steps

E.coli were harvested by centrifugation at 6,000 g, for 10 min, at 4°C (JLA 8.1000 rotor, Beckman), lysed in Lysis buffer (20 mM Hepes, 0.5 M NaCl, 2 mM MgCl2, pH7.2, 5% glycerol, 1 mM DTT) containing Protease Inhibitors EDTA-free (Bimake) and Benzonase (Merck), with a high pressure homogenizer LM-20 (Microfluidics), using two passages at 20,000 psi. Insoluble material was removed by high speed centrifugation (30,000g, 1 hour, at 4°C, in rotor JA12, Beckman) and by 0.45 *µ*m filtration. IMAC purification was performed with 5 ml HisTrap FF columns (Cytiva). After equilibration and loading, the column was washed with Lysis buffer, supplemented with 20 and then 50 mM imidazole. Finally, 10xHis-tagged-cCPE was eluted with IMAC elution buffer (20 mM Hepes, 0.5 M NaCl, pH7.2, 5% glycerol, 500 mM imidazole, 0.5 mM TCEP). To remove the 10xHis tag, the protein was incubated with HRV3C protease (Merck) while dialysed against a size exclusion buffer (20 mM Hepes pH7.2, 300 mM NaCl, 5% glycerol, 0.5 mM TCEP) overnight at 4°C. Size exclusion chromatography was performed with HiLoad Superdex200 column (Cytiva), equilibrated in 20 mM Hepes pH7.2, 300 mM NaCl, 5% glycerol, 0.5 mM TCEP.

#### 0.5.4 cCPE labelling with ATTO647

cCPE was labelled with ATTO647 maleimide (Sigma), according to the manufacturer’s protocol. Labelled cCPE was separated from free dye excess using first a desalting nap-5 column (Cytiva), followed by gel filtration, using a 24ml Superdex75 column (Cytiva) equilibrated in 20 mM Hepes pH7.2, 300 mM NaCl, 5% glycerol, 0.5mM TCEP. cCPE-labelled fractions were analysed by SDS PAGE. Positive fractions were pooled and protein concentrated. The degree of labelling was calculated to be 0.6.

### 0.6 Viscosity estimation with 3D particle tracking

To estimate viscosity in approx. 90% H_2_O and approx. 1% glycerol, Carboxyl Fluorescent Pink Particles (CF-2058-2, Spherotech, Inc.) was used at 1:10 dilution. To estimate viscosity in the lumen, spherical organoids were treated with Cell-Mask-orange (C10045, Invitrogen) at a concentration prior to image acquisition (Supplementary Figure 2a). Following approx. 2 hours of treatment CellMask-orange positive particles were imaged in a Spinning-disk microscope. A 3D stack imaging was performed with a z resolution between 0.25-0.35 µm and time-resolution of 0.7-1.2 sec depending on the size of the 3D stack at a given time point.

To obtain particle segmentation & trajectories, images were denoised by using Noise2Void [53]. Afterwards, particles were manually cropped in 3D and time. To segment the particles, a combination of StarDist and APOC (Accelerated Pixel and Object Classification) python-based tools were used to obtain 3D labels in time [54, 55]. To obtain particle trajectories, LapTrack & Napari-Laptrack were used to track particles according to their distance and image-based features (intensities and size) between time frames (Supplementary Figure 2b) [56].

For the estimate of the diffusion coefficients of the particles, trackpy was utilized to obtain 3D mean squared displacement curves from the particle trajectories obtained [57]. To obtain accurate diffusion coefficients, particles with tracks longer than 20 time frames and a regression of above 0.75 for the linear fit of the mean squared displacement were selected. To measure the size of particles, the biggest area (in the z-axis of the segmentation) was used to derive the radius of the particle. As a result, we observed that the particles had a negative correlation between particle radius and diffusion coefficients (Supplementary Figure 2e & f). To estimate the viscosity of fluids using the obtained particle radius and diffusion coefficients, we used the Stokes-Einstein equation (Supplementary Figure 2c).

### 0.7 Inference of hydrostatic lumen pressure with linear laser-ablation

First, the organoids were taken out of the Matrigel to perform the laser ablation. This was achieved by mechanically breaking the Matrigel dome (with organoids within) with pipettes followed by Liberase (CF-2058-2, Spherotech, Inc.) incubation at 37°C for 15 min to achieve enzymatic dissociation. Afterwards, individual organoids were transferred onto 8 well glass-bottom plates (80826,Ibidi) containing organogenesis medium w/o chemical treatments.

To create conduits accross the epithelium for inferring hydrostatic pressure of the lumen [12], a laser-ablation was performed by utilizing a Zeiss LSM 780 NLO system (Zen Black v11.00 software) with a 2-photon laser (Titatnium/Saphire). With the Zen Black software the 2-photon laser, with a power of 3.2 W at the laser head, was set to 100% laser power at a wavelength of 800 nm. A line scan with a width of 6.6-9.9 µm and 40–50-line repetitions across the epithelium at the middle plane of a lumen was performed to create a cut across the epithelium. Prior and post laser cutting, a 3D stack of the lumen was imaged (with a time resolution of 10-25 seconds: depending the size of the lumen) to later measure the lumen volume changes.

FiJi software was used to quantify the image-based variables for the Hagen-Poiseulle model. The line-tool and measure function was used to measure the monolayer thickness and radius of the channels (created by the laser ablation) [58]. LimeSeg, a Fiji plugin for the segmentation of 3D objects, was used to quantify the flow rate via lumen volumes changes before and after the laser-ablation [59].

### 0.8 Epithelial permeability assay

As a readout for epithelial permeability of pancreatic organoids, 3-5kDa Dextran-Alex488 (D22914, ThermoFisher Scientific) or 10kDa Dextran-Alex647 (D22914, ThermoFisher Scientific) were supplemented to the organoid media. After 1-3 hours of incubation, organoids were imaged with a spinning-disk microscope and light-sheet microscope.

### 0.9 Transcriptome Analysis

Branching and spherical organoids at day 7 of culture were lysed with lysis buffer RLT, and RNA was purified following the manufacturer instructions (RNeasy Plus Micro Kit, 74034, Qiagen). The quality of the purified RNA using an Agilent 2100 Bioanalyzer, following the instructions of the manufacturer (Agilent RNA 6000 Pico Kit 5067–1513). Amplification of the extracted RNA (700 pg) was performed by Ovation Pico SL WTA system V2 (3312–48, Nugen). Samples were labeled with SureTag DNA labeling kit (5190–3391, Agilent Technologies), run on SurePrint G3 Mouse Gene Exp v2 Array (G4852B, Agilent Technologies), and signals were read by a SureScan Microarray Scanner (Agilent Technologies).

### 0.10 EdU incorporation assay

Organoids were incubated with 10 *µ*M EdU (Click-iTPlus EdU Alexa Fluor 647; C10640, Invitrogen) in organogenesis medium for 2 hours at 37°C and 5% CO_2_. Then, organoids were processed for immunostaining as described above. Permeabilisation, blocking, and Click-iT reaction for EdU detection were performed according to the manufacturer’s instructions.

### 0.11 Microscopy

#### 0.11.1 Spinning-disk microscopy

All spinning-disk microscope images were performed with a Olympus IX 83 inverted stand driven by the Andor iQ 3.6 software. The microscope is equipped with a motorized xyz stage, a Yokogawa CSU-W1 scan head, and an Olympus 30 × 1.3 NA Sil Plan Apo objective. The setup was located inside a temperature-controlled chamber set to 37°C and 5% CO2. The sample was illuminated with a 561 nm laser, and the emission was collected using a 525/50 bandpass filter.

#### 0.11.2 Confocal microscopy

All confocal images were acquired using an inverted Zeiss LSM 780 confocal microscope with a 40×/1.3 NA C-Apo water objective, 561 nm diode laser, and a pinhole setting between 1 and 2 Airy Units. All images were acquired using Zen Black software (Carl Zeiss).

#### 0.11.3 Light-sheet microscopy

Organoids were grown in V-shaped bottom chambers made from Fluorinated ethylene propylene (FEP) from Viventis. A Viventis light-sheet microscope was used with a Nikon 25x 1.1 NA water immersion objective. The imaging parameters were set to: dual illumination of 2.2 *µ*m beams with Laser 561/647, and an exposure of 100 ms. The chamber was set to 37°C and 5% CO2. Organoids were imaged every 30 min intervals.

### 0.12 Image analysis and Quantification

#### 0.12.1 2D & 3D organoid and lumen segmentation and quantification

To segment the lumen and whole organoid structure, images were first denoised using Noise2Void [53]. Images containing epithelial markers (nuclei and membranes) were summed using pyclesperanto-prototype [60]. The summed epithelium channel was then processed with Gaussian blur (sigma for xyz axes = 0.75-1.5) and Top-hat background removal (radius for xyz axes = 20-30). These processed channels, along with the lumen within the epithelium, were manually annotated using Napari to create training data for an APOC model [54]. Using the trained APOC model, the epithelium channels were segmented. Inaccuracies in the prediction output were manually corrected with Napari or semi-automatically corrected using the binary processing functions of pyclesperanto-prototype. The lumen, identified as a 3D hole in the epithelium mask, and the segmentation output were used to generate triangulated meshes. To perform 2D segmentation of organoids and lumen structures, the largest area along the z axis was selected from the 3D segmentation output for further analysis and quantifications.

To generate meshes from the lumen and epithelium, 3D Marching-Cube function of scikit-image were applied on the lumen binary and the epithelium binary, that had been processed with the binary fill holes function of scipy-image and rescaled pixel of pyclesperanto-prototype for isotropic pixels [60, 61]. The generated meshes were smoothened using the Laplacian smoothening fucntion of Trimesh [62]. Other features of the lumen and organoid meshes were obtained via Trimesh functions: integrated mean curvature, volume, and surface area. To generate the topological skeleton of the lumen, the Skeletonizing function of the scikit-image were applied to the lumen binary [61].

The following calculations were performed to obtain the morphological features of the lumen and organoids:

- **Lumen and organoid sphericity**: to numerically characterize the 3D shape of the objects we quantified the reduced volume (*υ*) by applying the volume (*V*) and surface area (*SA*) obtained from the generated meshes (above) [15]. This quantification resulted in perfect spheres exhibiting a reduced volume of 1 and in lower values with decreasing sphericity.

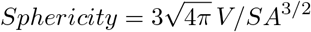
- **2D and 3D Lumen occupancy (*LO*)**: to obtain the 3D lumen occupancy, volumes (*V*) obtained from the 3D segmentation of the lumen and organoid we used. For 2D lumen occupancy, areas (*A*) obtained from the 3D segmentation of the lumen and organoid we used.

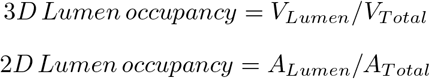

The lumen occupancy values are presented as percentages, except for Supplementary Figure 3.

#### 0.12.2 3D nuclei segmentation and quantification

The segmentation of nuclei in 3D-images was performed using StarDist [55]. First, a subset of images with nuclei staining were manually annotated using Napari as training data to create a StarDist model. After, the trained model was applied to predict and segment the nuclei. The nuclear segmented output was used to quantify the number of EdU-, DAPI-, and Hoescht-marked nuclei in organoids.

The following calculations were performed to obtain the proliferation features of the organoids:

- **EdU:DNA ratio**: To quantify active proliferation detected with the EdU incorporation assay we obtained the total number of EdU and DNA per organoid from the nuclear segmentation (above). From that we presented the data as ratio

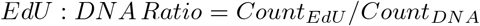
- **Doubling time**: To quantify the rate of cell population doubling we obtained the average number of cells at 48 hours 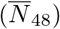 and 96 hours 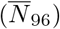

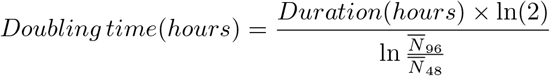

### 0.13 Mathematical model

#### 0.13.1 Phase field model

To computationally simulate the multicellular morphology, we applied the multicellular phase field model [63] with lumen phase [25, 30]. Although the details of the model used in this paper follow those in Tanida *et al*. [25], we changed the cell growth rule in the following way. In Tanida *et al*., we assumed the term *α*[*V*_*target*_ − *V*_*i*_(*t*)] in the free energy functional with the constant target volume *V*_*target*_ as well as the variable area of each (*i*-th) cell *V*_*i*_(*t*) and the constant prefactor *α*; in contrast, in this paper, this term is replaced by *α*[*V*_*target,i*_(*t*) − *V*_*i*_(*t*)] where *V*_*target,i*_(*t*) is the time-dependent target volume, which allows us to control the cell growth rule. The time evolution of *V*_*target,i*_(*t*) was assumed to obey

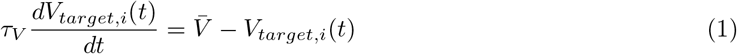

(*i.e*. exponential convergence toward the given constant 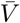 with the given characteristic convergence time *τ*_*V*_), so that we can control the typical cell growth duration by tuning the parameter *τ*_*V*_. Finally, in this paper, we do not assume the minimum duration for the division of each cell after the division of its mother cell, which we assumed in Tanida *et al*. Parameter values used in this paper are as follows; *α* = 1.0, 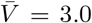, with *τ*_*V*_ = 1, 10, 20, 30, 40, 50, 60, 70, 80, 90 and *ξ* varying from 0.10 to 0.32 with increments of 0.02. In the model, at the end of cell division, micro-lumens are created at the middle point of the spindle poles (of a dividing cell) with a fixed size of value 0.7. The other parameter values and the initial conditions are set identical to those of Tanida *et al*.

#### 0.13.2 Relationship between *ξ* and Δ*P* in the phase field model

In the Phase Filed Model (PFM), the relationship between *ξ* and Δ*P* has been derived analytically in Tanida *et al*. from the growth dynamics of the domain in the two-dimensional (2D) case [25]. Considering the growth of the isolated lumen in one dimension first, the free energy associated with the lumen variable *u*_0_ in one dimension is integrated by using the analytic solution of the interface, 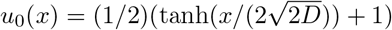, as

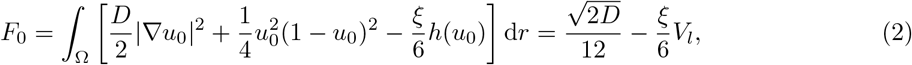

where *D* is the diffusion coefficient, *h*(*u*) = *u*^2^(3 − 2*u*), and *V*_*l*_ = ∫_Ω_ *h*(*u*_0_)d*r* is the volume of the lumen. The relation is extended to a 2D case and the growth of lumen volume *V*_*l*_ was derived from the kinetic equation based on the gradient flow of the free energy as,

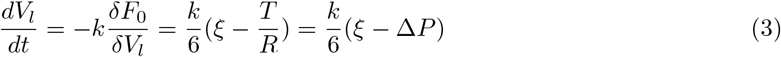

where *k* is the kinetic constant, 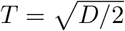 is the tension of the interface, and 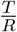 can be regarded as the pressure difference, Δ*P*, between the inside and outside of the cyst in analogy to the lumen growth kinetics equation [64], This relation was confirmed successfully by numerical simulation.

The dynamics of lumen growth are described by the following equation [64],

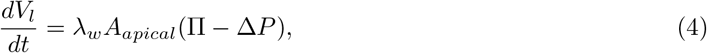

where *V*_*l*_ is the lumen volume, *λ*_*w*_ is the water permeability of the cell layer, *A*_*apical*_ is the apical area of the lumen surrounding the cell layer, Π is the osmotic pressure difference between the inside and outside of the cyst, Δ*P* is the hydrostatic pressure difference between the inside and outside. This relation can be written by using the lumen radius *R* for spherical lumen. Applying the relation *V*_*l*_ = *πR*^2^ and *A*_*apical*_ = 2*πR* in 2D, Eq. (2) becomes

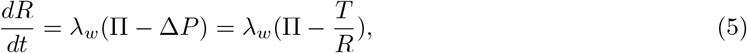

where 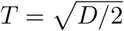. For 3D case, employing 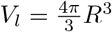 and *A*_*apical*_ = 4*πR*^2^ leads to

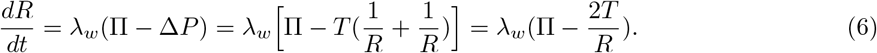

If a cell layer surrounds the lumen, the tension in the cell layer is expected to add to T. In such a case, total tension becomes *T*_*total*_ = *T* + *T*_*cell*_, where *T*_*cell*_ denotes the tension by the cell layer.

To account for the effect of cell layer tension in PFM, a two-dimensional simulation of a cyst surrounded by a cell layer was performed with a parameter set and with the same rule in the simulation performed. By varying *ξ*, we obtain the time series of *V*_*l*_(*t*). Assuming that the lumen is spherical, we obtain 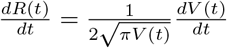. Then, Δ*P* can be estimated as

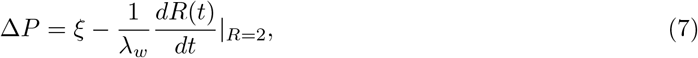

where 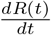 depends on the value of *R*, therefore we used the value at *R* = 2. Estimated Δ*P* values from the time series of *V*_*l*_(*t*) and *R*(*t*) for various *ξ* are shown in Supplementary Figure 4. From the best fit in Supplementary Figure 4, we obtain Δ*P* = 0.89*ξ* 0.0068. The slope is slightly less than 1, with an offset corresponding to the presence of a critical osmotic pressure required for lumen growth.

Note that this is a two-dimensional case; in the three-dimensional case, the three-dimensional hydrostatic pressure difference, denoted by Δ*P*_3_, is twice larger than the two-dimensional value for the same tension *T*. Thus, we employ the relation, Δ*P*_3_ = 2Δ*P*_2_, where Δ*P*_2_ denotes the hydrostatic pressure difference in 2D.

This yields an empirical relationship between *ξ* and Δ*P* in the 3-dimensional PFM, Δ*P*_3_ = 2 *×* (0.89*ξ* − 0.068) = 1.78*ξ* − 0.136. Thus, if the ratio of measured hydrostatic pressure difference between the sphere and pancreas organoid is Δ*P*_*sphere*_ = 3.4Δ*P*_*org*_, then the ratio of the osmotic pressure between sphere and organoid is *ξ*_*sphere*_ = 3.4*/*1.78 × *ξ*_*org*_ = 1.9 × *ξ*_*org*_. We use this conversion relationship between Δ*P* and *ξ*, which is expected in the three-dimensional case in the PFM, in our comparison with the experimental data.

## Material and code availability

Material and image analysis code requests should be addressed to the corresponding author. The software code used for the phase field model simulation is available in the Github: https://github.com/kana-fuji/MCPFM_tauV-model.git

## Acknowledgments

We extend our gratitude to Dr. Alf Honigmann, Dr. Markus Murkenhirn, Dr. Linjie Lu, and Dr. Tristan Guyomar for their invaluable discussions as part of the HFSP team. Special thanks to Dr. Sakurako Tanida for her contribution to the phase field modeling. We are deeply indebted to Congtin Justin Nguyen for his pivotal role in establishing the permeability assay, and to Robert Haase at the Data Science Center ScaDS.AI, University of Leipzig, for his support and guidance in image analysis.

Additionally, we are grateful to Barbara Borgonovo, Aliona Bogdanova, and Eric Geertsma of the Protein Biochemistry Facility at MPI-CBG for their assistance in synthesizing the cCPE-647 toxin. We also thank Riccardo Maraspini, Catarina Nabais, Britta Schroth-Diez, and Jan Peychl from the Light Microscope Facility at MPI-CBG for their invaluable support in imaging. Further acknowledgments go to Josefin Krull and Petra Huebner from the Biomedical Facility at MPI-CBG for their dedicated services in mouse maintenance.

Byung Ho Lee was supported by the Early Postdoc Mobility Swiss National Science Foundation grant (project number: P2GEP3-181529). This project was funded by HFSP project number RGP0050/2018.

## Author contributions

Conceptualization and funding, B.H.L., K.F., D.R., T.H., M.S. and A.G.-B.; Experimental Investigation, B.H.L., H.P., P.S., S.Y. and C.S.; Data Analysis B.H.L., H.P., S.Y. and A.L.; Theoretical and computational modelling K.F., T.H. and M.S.; Writing B.H.L.and A.G.-B. with inputs from all authors; All authors discussed the results and commented on the manuscript.

