## Supplementary Material for "Control of lumen geometry and topology by the interplay between pressure and cell proliferation rate in pancreatic organoids"

### Supporting Information

#### Figure S1

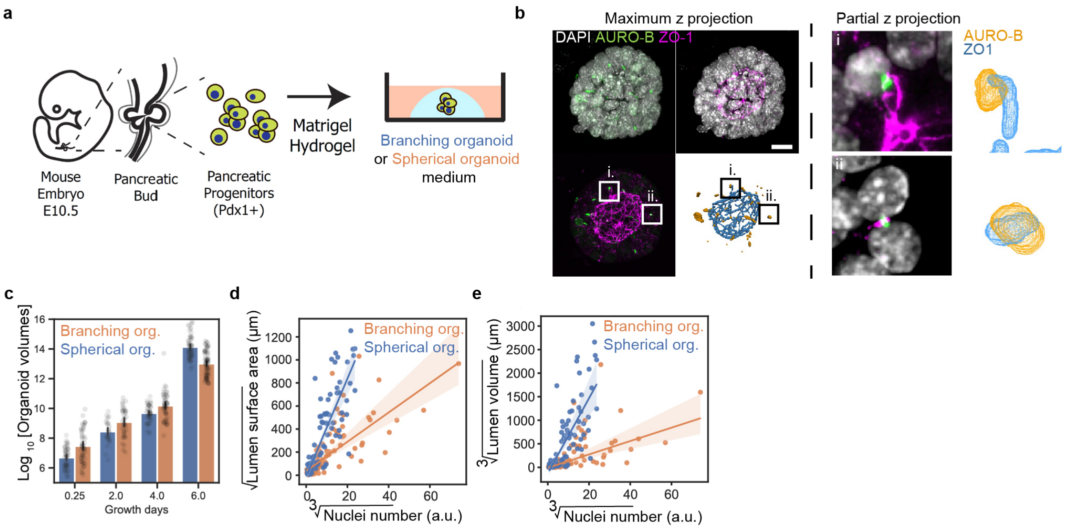


**Figure S1: Initiation and evolution of lumen in pancreatic branching organoids and spherical organoids. (a)** Schematic of pancreatic organoid growth protocol. **(b)** Left: Maximum projected immunofluorescence images of spherical organoids stained with DAPI (gray, nuclei), Aurora-B (green, abscission point), ZO1 (magenta, sub-apical tight junctions). In addition, mesh representation of segmented Aurora-B and ZO1. Right: Partial z-projected zoomed-in images, showing a co-localization of Aurora-B and ZO1. Scale bar: 10 µm. **(c)** Quantification of organoid volume (expressed as $\log_{10}$[organoid volume] of branching and spherical organoids. N=3-5 exp; n=220 branching organoids, 194 spherical organoids. **(d)** Quantification showing the positive correlation between lumen surface area and nuclei number for branching and spherical organoids. N=3-5, n=54 branching organoids, n=72 spherical organoids. **(e)** Quantification showing the positive correlation between lumen volume and nuclei number for branching and spherical organoids. N=3-5, n=54 branching organoids, n=72 spherical organoids.

### Figure S2

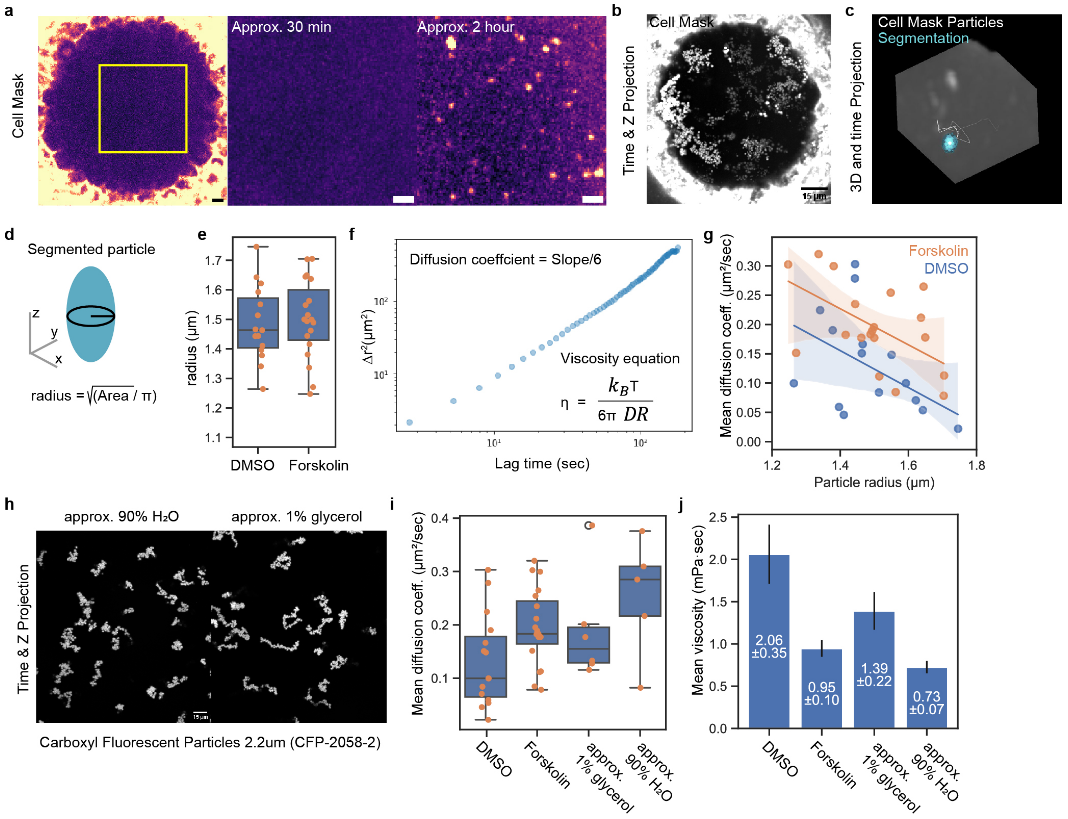


**Figure S2: Estimation of lumen viscosity for lumen-to-exterior hydrostatic pressure inference.** **(a)** Live images of spherical organoids at approximately 30 min and 2 hours Cell Mask treatment. Scale bar=5 µm. **(b)** Time- and z-projection image of Cell Mask-treated spherical organoid. Scale bar=15 µm. **(c)** 3D projection and trajectory of segmented Cell Mask-positive particle. **(d)** Schematic representing the measurement of radius from the 3D segmented particle. **(e)** Quantification of Cell Mask-positive particles radius in DMSO- and Forskolin-treated spherical organoids. n = 15 DMSO, 19 Forskolin. **(f)** An example of a 3D Mean squared displacement curve of a Cell Mask-positive particle tracked over time. Insert shows equations for Diffusion Coefficient ($D$) and fluid viscosity ($\eta$). **(g)** Quantification of diffusion coefficient of particles with various sizes (represented as radius) that were tracked in DMSO- and Forskolin-treated spherical organoids. n = 15 DMSO, 19 Forskolin. Linear-fit plot with 95% confidence interval error bars. **(h)** Time- and z-projected images Carboxyl Fluorescent Particles (2.2µm diameter) in approx. 90% $H_2O$, and approximately 1% glycerol. Scale bar = 15 µm. **(i)** Quantification of Diffusion coefficient of particles tracking in DMSO- and Forskolin-treated spherical organoid lumens,approx. 90% $H_2O$, and approx. 1% glycerol. n = 15 DMSO, 19 Forskolin, 6 approx. 1% glycerol, 4 approx. 90% $H_2O$. **(j)** Quantification of fluid viscosity of DMSO- and Forskolin-treated spherical organoid lumens and in approx. 90% $H_2O$, and approximately 1% glycerol solutions. n = 15 DMSO, 19 Forskolin, 6 approx. 1% glycerol, 4 approx. 90% $H_2O$.

### Figure S3

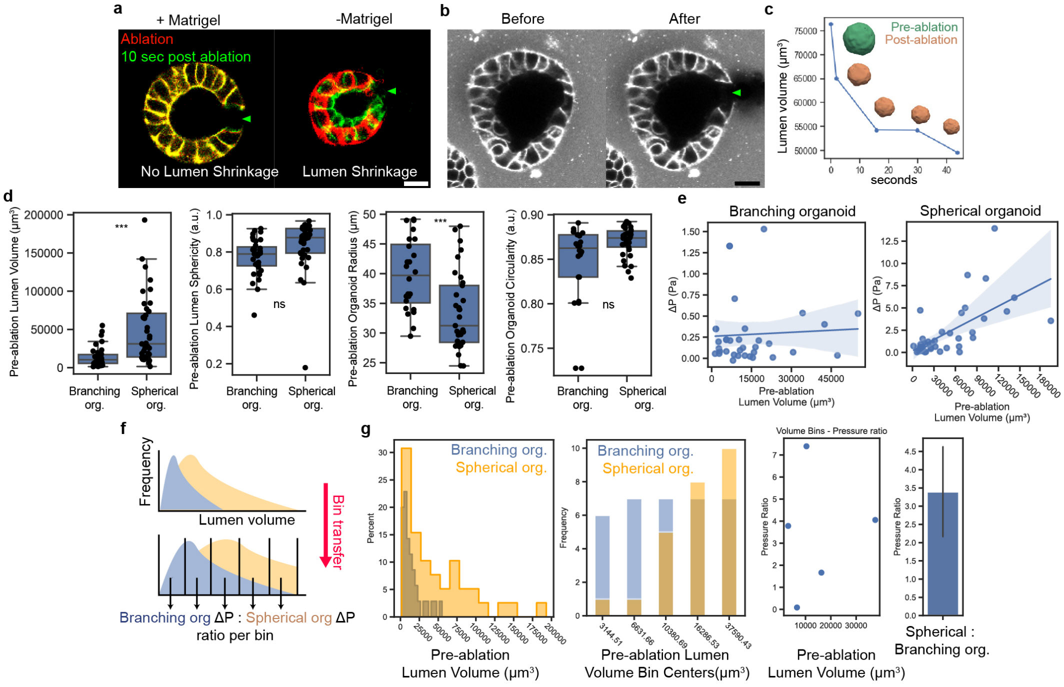


**Figure S3: Lumen-to-exterior hydrostatic pressure inference of branching and spherical organoids.** **(a)** Live images of laser-ablated spherical organoids in Matrigel with no lumen shrinkage (left) and removed from Matrigel with lumen shrinkage (right). Green arrowheads: conduits created by laser ablation. **(b)** Cell Mask-treated spherical organoids before and after laser ablation. Green arrowheads: conduits created by laser ablation. **(c)** Quantification of lumen volume over various acquisition time points during laser ablation experiment. Insert show 3D meshes of the segmented lumen exemplifying the change morphology and size of the lumen before (green) and after (beige) laser ablation. **(d)** 3D Quantification of lumen volume, sphericity, organoid radius, and organoid circularity before laser ablation. N = 3 exp; n = 26 branching organoids, 29 spherical organoids. **(e)** Quantification of lumen-to-exterior hydrostatic pressure with various volumes before laser ablation (left: branching organoids, right: spherical organoids). N = 3, n = 36 branching organoids, 34 spherical organoids. Linear-fit plot with 95% confidence interval error bars. **(f)** Schematic representation showing how comparative lumen-to-exterior hydrostatic pressure ratios relative to lumen volume are obtained. **(g)** Left: distribution of lumen volume in branching (blue) and spherical organoids (yellow). Mid: Bin transfer from branching organoids to spherical organoids. right: Mean lumen-to-exterior hydrostatic pressure ratios between branching and spherical organoids per bin. The Mann-Whitney test was performed for all statistical significance tests.

#### Figure S4

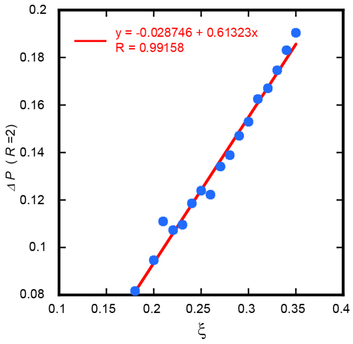


**Figure S4: Relationship between $\xi$ and $\Delta$P.** Estimated $\Delta P$ as a function of $\xi$ from 2D simulation of cysts with varying $\xi$, where the values of $dR(t)/dt$ were evaluated at $R = 2$.

#### Figure S5

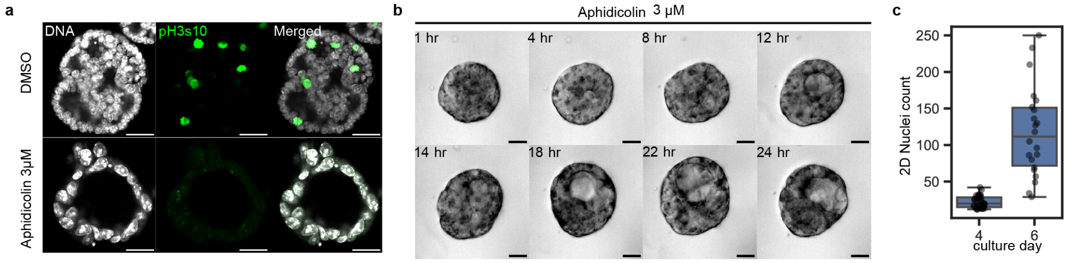


**Figure S5: *in vitro* and *in silico* Lumen morphogenesis under proliferation-arrested conditions. (a)** Immunofluorescence images of branching organoids treated with DMSO and Aphidicolin. Green: phospho-Histone 3 serine 10 (pH3s10), white: DNA. Scale bar = 30 µm **(b)** Montage of live imaging of Aphidicolin treated branching organoids. Scale bar = 20 µm. **(c)** Number of nuclei counted in the mid-plane (2D counting) of branching organoids at day 4 and day 6 of culture. n= 35 day 4, 22 day 6.

#### Figure S6

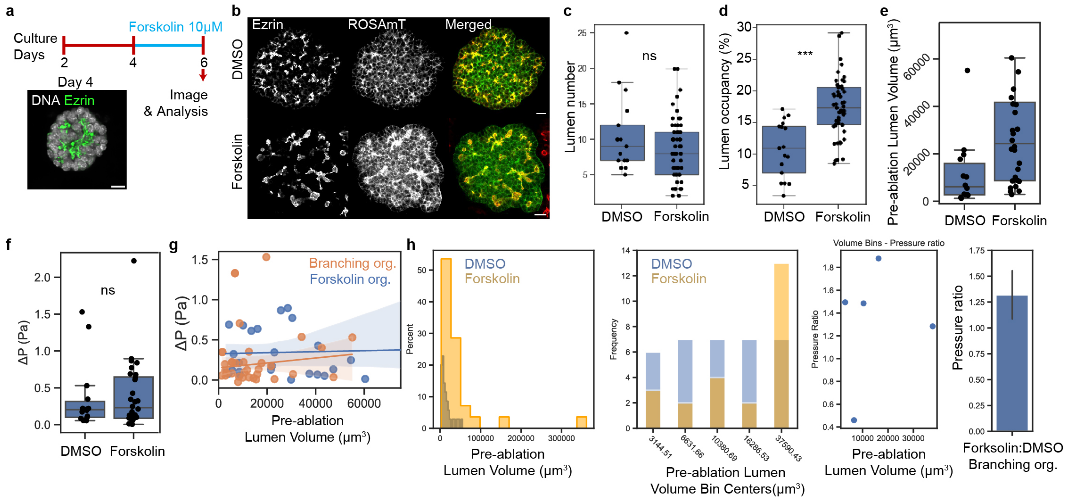


**Figure S6: Lumen-to-exterior hydrostatic pressure inference of branching organoids under Forskolin treatment.** **(a)** Schematic showing experimental design of forskolin treatment. **(b)** Immunofluorescence image of DMSO and forskolin treated branching organoids at day 6. Scale bar = 20 μm. **(c)** 2D Quantification of lumen number of DMSO and Forskolin treated branching organoids at day 6. N = 3 exp, n = 27 DMSO, 31 Forskolin. **(d)** 2D Quantification of lumen occupancy of DMSO and Forskolin treated branching organoids at day 6. N = 3 exp, n = 27 DMSO, 31 Forskolin. **(e)** 3D Quantification of lumen volume in branching organoids treated with DMSO and Forskolin before laser ablation N=3, n= 26 DMSO, 29 Forskolin. **(f)** Inference of lumen-to-exterior hydrostatic pressure $\Delta P$ of the DMSO and Forskolin treated branching organoid lumens on day 6 of culture. N = 3 exp; n = 26 DMSO, 29 Forskolin. **(g)** Quantification showing the relationship between lumen-to-exterior hydrostatic pressure and pre-ablation lumen volume of branching organoids under DMSO and Forskolin treatment. N=2-3 exp; n=26 DMSO, 29 Forskolin condition. **(h)** Left: Distribution of lumen volume in branching organoids (blue) and Forskolin-treated branching organoids (yellow). Mid: Bin transfer from branching organoids to Forskolin-treated branching organoids. Right: Mean lumen-to-exterior hydrostatic pressure ratios between branching and spherical organoids per bin. And the lumen-to-exterior hydrostatic pressure ratios of forskolin: DMSO treated branching organoids. Error bar: standard error of the mean.

### Figure S7

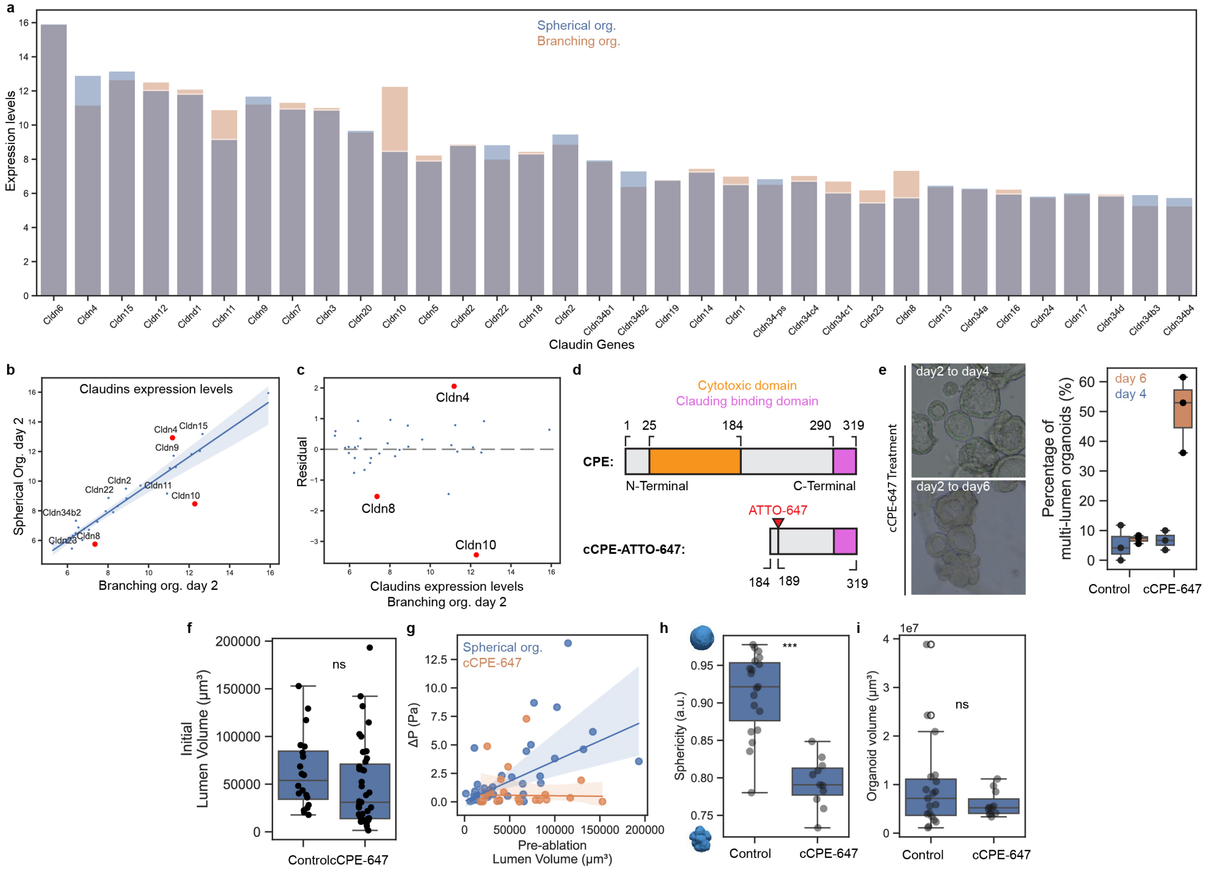


**Figure S7: Lumen morphogenesis under permeabilization with cCPE-647.** **(a)** Microarray transcriptomics data showing claudin expression levels (in log2) in branching organoids (blue) and spherical organoids (orange) on day 2 of culture. **(b)** Scatter Plot with Linear Regression showing the relationship between expression levels (in log2)of claudin genes in branching organoids and and spherical organoids at day 2 of culture. Top 10 genes that are deviating from the linear regression are annotated. Red points representing the top 3 deviating genes. **(c)** Scatter plot showing the residuals of the linear regression model (from b). A gray dashed horizontal line at y=0 indicates where residuals would lie if there were no deviation from the model. **(d)** Schematic diagram showing the full CPE (top) and truncated cCPE (bottom) linear structures, along with their cytotoxic and claudin-binding domains, and the location of ATTO-647. **(e)** Left: Phase-contrast images of spherical organoids treated with cCPE-647 from day 2 to day 4 (left) and day 2 to day 6 (right) of culture growth. Right:Quantification of the percentage of organoids showing multi-lumen phenotype in control and cCPE-647 treated spherical organoids from day 2 and analyzed at day 4 and 6. N=3 exp. **(f)** Quantification of pre-ablation lumen volume of spherical organoids under control and cCPE-647 treatment condition. N=2-3 exp; n=39 control, 20 cCPE-647. **(g)**Quantification showing the relationship between lumen-to-exterior hydrostatic pressure and pre-ablation lumen volume of spherical organoids under control and cCPE-647 treatment. N=2-3 exp; n=39 control, 20 cCPE-647. **(h-i)** 3D quantification of sphericity (f) and organoid volume (g) of control and cCPE-647 treated spherical organoids on day 6. N = 2-3 exp; n = 27 control, 17 cCPE-647. The Mann-Whitney test was performed for all statistical significance tests.

#### Supplementary Video 1

**Video S1: Cell and lumen dynamics in *in silico* organoid.** Growth of *in silico* organoids with $\tau_V = 20$ and $\xi = 0.26$. Red regions represents the cells. Blue regions represents lumen.

#### Supplementary Video 2

**Video S2: Cell and lumen dynamics in *in silico* organoid with no stop in cell division.** Growth of *in silico* organoids with $\tau_V = 40$ and $\xi = 0.12$. Red regions represents the cells. Blue regions represents lumen.

#### Supplementary Video 3

**Video S3: Cell and lumen dynamics in *in silico* organoid with stop in cell division.** Growth of *in silico* organoids with $\tau_V = 40$ and $\xi = 0.12$. Cell division was stopped when *in silico* organoids reached 32 cell numbers. Red regions represents the cells. Blue regions represents lumen.
